# Restored tactile sensation improves neuroprosthetic arm control

**DOI:** 10.1101/653428

**Authors:** Sharlene N Flesher, John E Downey, Jeffrey M Weiss, Christopher L Hughes, Angelica J Herrera, Elizabeth C Tyler-Kabara, Michael L Boninger, Jennifer L Collinger, Robert A Gaunt

## Abstract

The sense of touch is critical for skillful hand control^1–3^, but is largely missing for people who use prosthetic devices. Instead, prosthesis users rely heavily on visual feedback, even though state transitions that are necessary to skillfully interact with objects, such as object contact, are relayed more precisely through tactile feedback^4–6^. Here we show that restoring tactile sensory feedback, through intracortical microstimulation of the somatosensory cortex^7^, enables a person with a bidirectional intracortical brain-computer interface to improve their performance on functional object transport tasks completed with a neurally-controlled prosthetic limb. The participant had full visual feedback and had practiced the task for approximately two years prior to these experiments. Nevertheless, successful trial times on a commonly used clinical upper limb assessment task were reduced from a median time of 20.9 s (13.1 - 40.5 s interquartile range) to 10.2 s (5.4 - 18.1 s interquartile range) when vision was supplemented with microstimulation-evoked cutaneous percepts that were referred to different fingers and were graded in intensity based on real-time prosthesis contact forces. Faster completion times were primarily due to a reduction in the amount of time spent attempting to grasp objects. These results demonstrate the importance of tactile sensations in upper-limb control and the utility of creating bidirectional brain-computer interfaces to restore this stream of information using intracortical microstimulation.

---

We use our hands to interact with our environment, often by exploring and manipulating objects. Without tactile somatosensory feedback, even simple manipulation tasks become clumsy and slow^1–3^. Outside of investigational settings, this source of feedback is rarely provided for prosthetic devices^8^, and in the context of human brain-computer interfaces (BCIs), has only recently become possible^7,9–11^. These studies have begun to describe the perceptual characteristics of cortical stimulation, however, the potential benefits of a bidirectional BCI on function have remained unexplored. This is despite the fact that the need for somatosensory feedback in BCIs has long been suggested as the next step towards complete upper-limb restoration^12–14^ and cited by amputees as a desired feature^15–17^. Here we show that a bidirectional BCI (Fig. 1) that provides these tactile percepts improves performance in functional object transport tasks using a BCI-controlled robotic arm. The percepts were driven in real-time by sensors in a prosthetic hand (Fig. 1c,d), evoked through intracortical microstimulation (ICMS) of area 1 of somatosensory cortex (S1) and experienced by a participant as originating from his own palm and fingers.

**Fig. 1:**
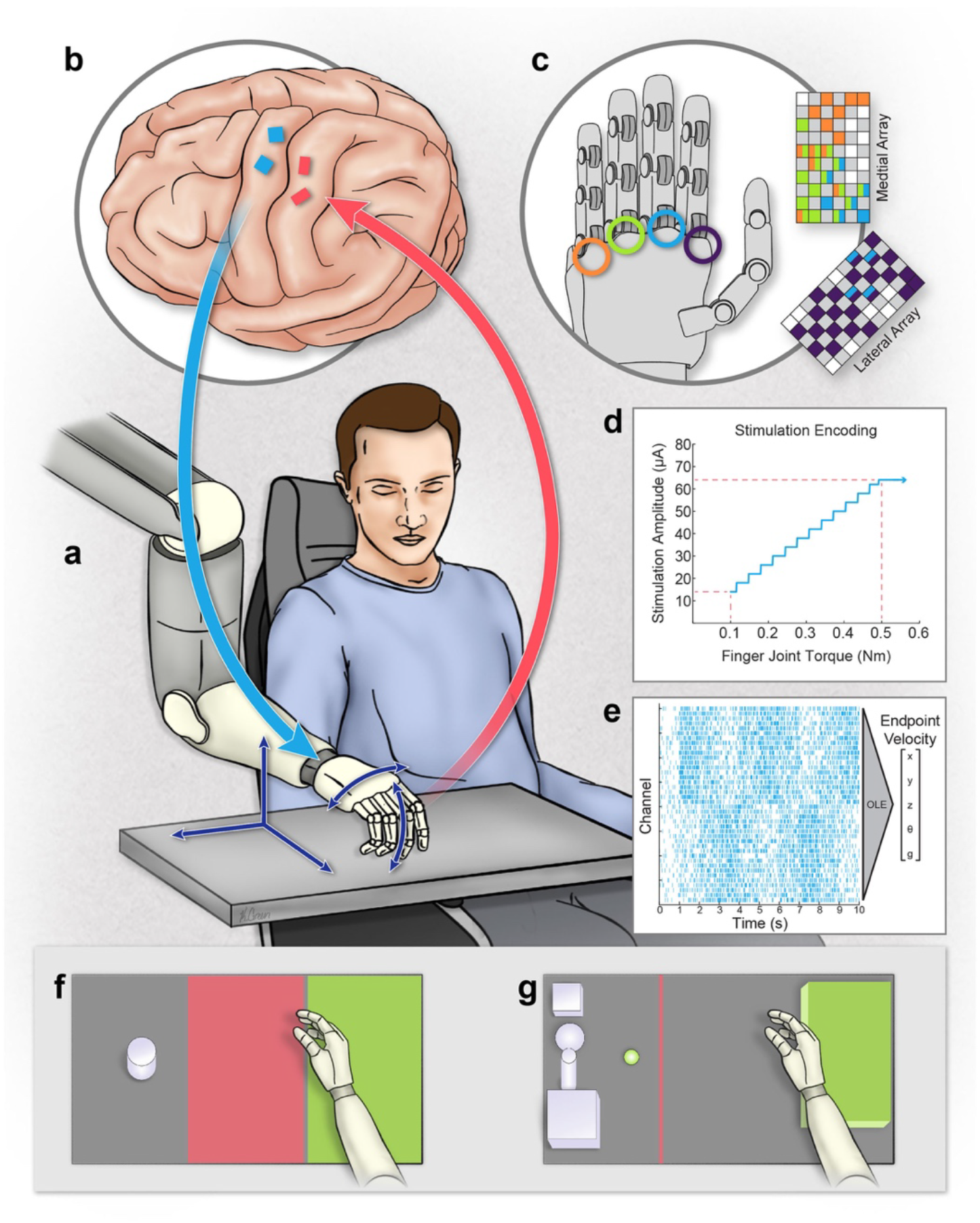
Overview of bidirectional BCI system components, operation and tasks. **a**, The study participant used a bidirectional intracortical BCI to control a robotic prosthesis in real time. The arm was positioned near the participant to provide clear visual feedback, but physical contact was not possible. The participant controlled the prosthesis in five dimensions, illustrated by the dark blue arrows (3D translation, wrist rotation and grasp). **b**, Four microelectrode arrays were implanted in the left hemisphere. Arrays in primary motor cortex (blue) recorded signals which were used to control the modular prosthetic limb. Arrays in somatosensory cortex (red) delivered stimulation pulses, which artificially activated neurons, resulting in sensory percepts referred to the hand. **c**, Stimulation of the electrode arrays in the somatosensory cortex evoked percepts from the base of the fingers. Colored grids represent individual electrodes on the two microelectrode array and the locations on the hand where stimulation through each electrode evoked a percept (index finger = purple, middle finger = blue, ring finger = green, little finger = orange)^7^. Torque sensors in the robot fingers were used to drive selected electrodes in the somatosensory cortex with matching somatotopic fields (e.g. index finger torque sensor controlled electrodes evoking percepts in the index finger). **d**, The torque measured at the base of the fingers increased as more force was applied to the objects. Stimulation current amplitude was modulated by torque using a linear transformation. **e**, Threshold crossing events were detected from the multichannel neural recordings in the motor cortex. Each row represents an individual electrode and each mark represents a threshold crossing event. Using an optimal linear estimation decoding scheme, endpoint velocity (v_x_, v_y_ v_z_) as well as wrist pro/supination velocity (v_θ_) and grasp velocity (v_g_) were simultaneously and continuous decoded. **f**, Overhead view of the object transfer task showing the grasp (gray area), transport (red area) and release (green area) zones. The cylindrical object was placed in the grasp zone by the experimenter, was grasped using the prosthesis, moved over the transport zone and placed in the release zone. This process was repeated as many times as possible in two minutes. **g**, Overhead view of the Action Research Arm Test (ARAT) showing the object presentation position (green dot) and the raised platform target (green box). Different objects (not all objects shown) were positioned at a standard location, grasped and then placed on the platform as quickly as possible. For all tasks, the arm was under full control of the user from the start to the end of a trial.

We used two tasks to evaluate performance: an object transfer task (Fig. 1f) and a modified version of the Action Research Arm Test (ARAT)^18^ (Fig. 1g). Both tasks were completed using the Modular Prosthetic Limb (MPL)^19^. The robotic arm was controlled using neural activity recorded from two 88-channel microelectrode arrays implanted chronically in primary motor cortex (M1) (Fig. 1b) of a human participant with tetraplegia resulting from a cervical spinal cord injury. Five degrees-of-freedom (DoF), consisting of 3D endpoint translation, pronation/supination of the wrist, and hand grasp aperture (Fig. 1a)–with the hand in a power grasp conformation–were continuously and simultaneously controlled by the participant during all tasks (Fig. 1e). Tactile feedback was delivered in the first four experimental sessions by ICMS through two 32-channel microelectrode arrays implanted in area 1 of S1 (Fig. 1b). Stimulation pulses were delivered at 100 pulses per second and pulse amplitude was modulated linearly by the reaction torques measured at the metacarpophalangeal joint of the fingers on the MPL (Fig. 1d). Pulse trains were delivered to electrodes which, when stimulated, evoked percepts on corresponding fingers (Fig. 1c).

We first tested the effect of providing ICMS-induced tactile feedback on functional performance using an object transfer task that was familiar to the participant. The goal was to transport a compliant object across the workspace (Fig. 1f) as many times as possible in two minutes (Supplemental Video 1). We compared the number of transfers completed during four sessions with ICMS to four sessions without ICMS. Each session consisted of five two-minute trials. Across a total of 20 trials with ICMS, 352 transfers were completed compared to 315 transfers in the 20 trials without ICMS (Table 1). The number of transfers increased from 15.8 ± 3.8 transfers per trial to 17.8 ± 2.4 transfers per trial with ICMS, though this difference was not statistically significant (t_38_ = −2.02, P = 0.050, *t*-test). However, we observed qualitative improvements during the task that led us to examine the data in more detail.

**Table 1:** Performance metrics for each task per experiment day. The total number of object transfers is the sum of all five 2-minute trials per day. ARAT scores were computed as the sum of the best score per object, with a maximum score of 27. Each of the nine objects was attempted 3 times, so that the maximum number of trials attempted per session was 27. The total median and IQR trial time for successful ARAT trials was calculated by pooling trial times across all four sessions per feedback condition and calculating the median and IQR from the aggregate distribution.

|  | Session | Object Transfer<br>(transfers per<br>day) | ARAT Score<br>(out of 27) | ARAT Trials Completed<br>(out of 27) | Median and IQR trial<br>time for Successful<br>ARAT Trials (s) | Median Sequence<br>Task % Correct |
| --- | --- | --- | --- | --- | --- | --- |
| With ICMS<br>Feedback | 1 | 97 | 21 | 19 | 11.9 (6.6 – 27.7) | 90 |
|  | 2 | 74 | 21 | 22 | 12.0 (5.6 – 38.9) | 90 |
|  | 3 | 93 | 21 | 21 | 8.8 (6.0 – 17.2) | 100 |
|  | 4 | 88 | 20 | 19 | 8.1 (4.6 – 11.9) | 100 |
|  | **Total** | **352** | **83** | **81** | **10.2 (5.4 – 18.1)** |  |
| Without ICMS<br>Feedback | 1 | 88 | 19 | 23 | 14.0 (11.1 – 30.9) | 100 |
|  | 2 | 55 | 16 | 19 | 27.6 (18.8 – 37.2) | 100 |
|  | 3 | 74 | 17 | 23 | 18.7 (12.3 – 41.7) | 100 |
|  | 4 | 98 | 17 | 13 | 40.5 (15.5 – 48.4) | 100 |
|  | **Total** | **315** | **69** | **78** | **20.9 (13.1 – 40.5)** |  |

The object transfer task can be broken up into grasp, transport and release phases. We defined these phases using the physical location of the MPL hand. The transport zone consisted of a region 22.5 cm wide and centered on the starting location of the hand at the beginning of a trial. The grasp zone was located to the left side of the transport zone, while the release zone was located to the right (Fig. 1f). We first examined the amount of time spent in each movement zone per transfer. We found that the time spent in the grasp zone decreased from 3.3 ± 1.2 s per transfer without ICMS to 2.3 ± 0.4 s per transfer with ICMS (t_38_ = 3.3, P = 0.002, *t*-test, Fig. 2a) while time spent in the release zone decreased from 2.8 ± 1.0 s per transfer without ICMS to 2.3 ± 0.5 s per transfer with ICMS (t38 = 2.0, P = 0.048, *t*-test, Fig. 2a). Time spent in the transport zone per transfer was no different with or without ICMS (2.1 ± 0.6 s without ICMS, 2.3 ± 0.3 s with ICMS, t_38_ = −1.3, P = 0.206, *t*-test, Fig. 2a). To uncover the reason behind the lower grasp times with ICMS, we examined the total distance travelled while the MPL was in the grasp zone. We found that there was significantly more movement in the grasp zone in trials without ICMS compared to trials with ICMS (44.2 ± 13.1 cm/transfer without ICMS, 32.4 ± 5.9 cm/transfer with ICMS, t_38_ = 3.7, P = 0.0007, *t*-test, Fig. 2b). This suggests that in trials without ICMS, the additional time was used to move the hand into an ideal configuration to grasp the object. This effect is further illustrated by comparing the spatial distributions of time spent across the workspace per transfer (Fig. 2c). With ICMS-evoked sensations, the participant spent less time in the immediate vicinity of the object.

**Fig. 2:**
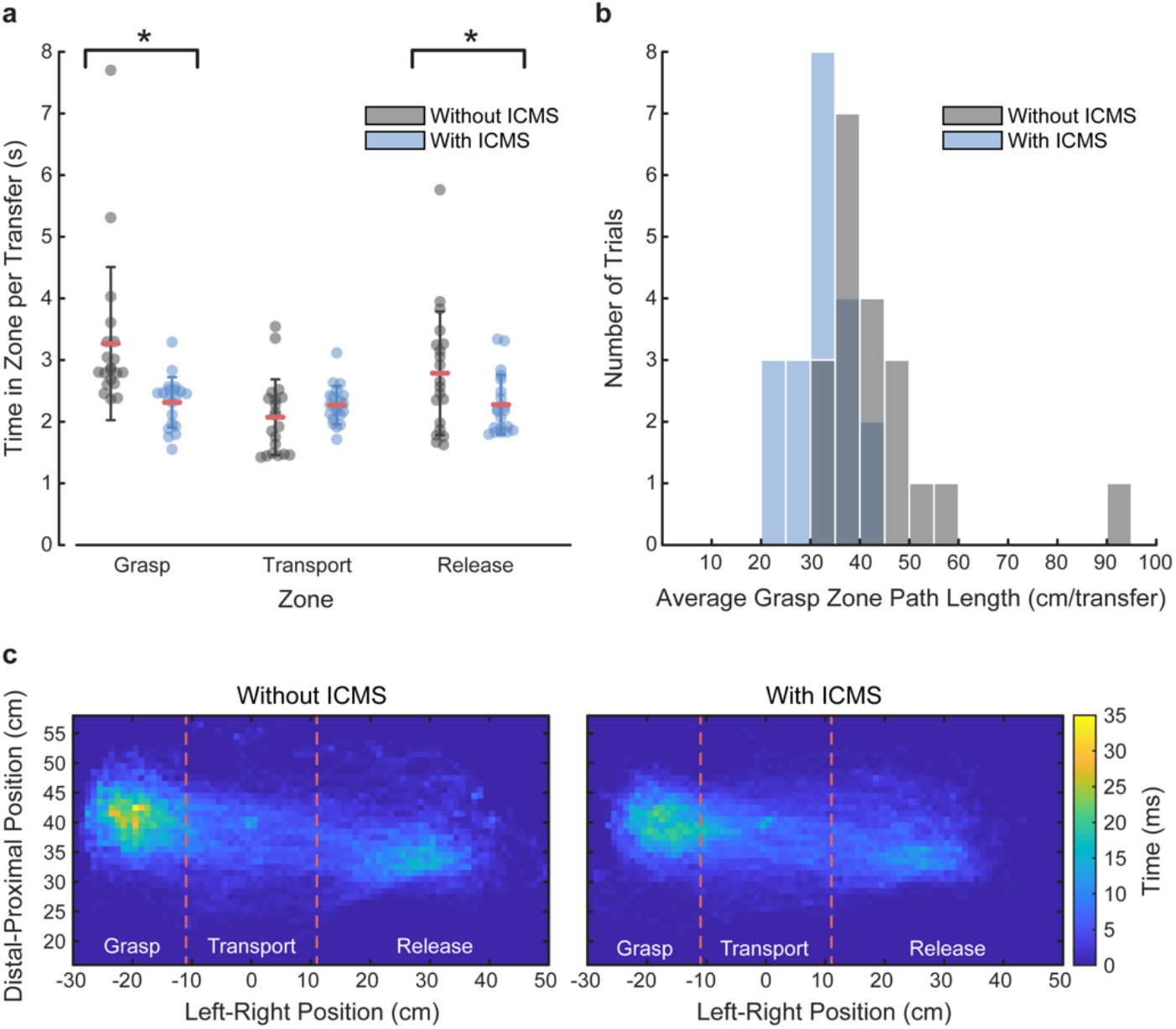
Object transfer performance. **a**, Amount of time spent in each task zone, per transfer, by feedback condition (n = 20 trials per feedback condition). Data for all trials are shown with red lines indicating the mean value and the whiskers indicating one standard deviation. The amount of time spent in the grasp and release zones decreased significantly with ICMS feedback (*P = 0.002 and 0.048, *t*-test, respectively), but the amount of time in the transport zone per transfer was not affected. **b**, Distribution of average path lengths in the grasp zone per trial for the two feedback conditions, computed as the total path length divided by the number of transfers. The longer path lengths (*P = 0.0007, *t*-test) without ICMS suggest that the extra time spent in the grasp zone was to adjust the endpoint position, rather than to hold the robot still while attempting to issue a grasp command. **c**, Spatial map of the amount of time spent in each location in the workspace per transfer. Each individual square represents a 1 x 1 cm region. Without stimulation, there was substantially more time spent near the object in the grasp zone as shown by the increase in the number of locations colored yellow in the grasp zone. Red lines indicate zone boundaries. Color indicates the amount of time spent in each location per transfer.

We then compared performance on a modified version of the ARAT^18^, which is a clinically validated test of unilateral upper-limb function and one that has been used previously to assess arm control performance in BCI systems^20,21^. We placed different objects on the left side of the workspace, one at a time, and asked the participant to grasp the object and place it on a raised platform on the right side of the table as quickly as possible (Fig. 1g and Supplemental Videos 2-4). A score of three was awarded if the task was completed in under five seconds, a score of two was awarded if the task was completed in under two minutes and a score of one was awarded if the object was touched but the task was not completed in two minutes. A score of zero was awarded otherwise. Each of the nine objects were attempted three times, for a total of 27 trials per ARAT session. The final score was the sum of the best score of the three attempts for each object.

Prior to these experiments, the participant had performed 23 ARAT sessions over a period of 23 months using several different control schemes, including four preliminary sessions with ICMS-driven tactile feedback (Fig. 3a). These four exploratory sessions included ICMS, but did not have consistent mapping between finger torque feedback and stimulation parameters. Further, these sessions were intermixed with sessions without ICMS rather than being performed consecutively with fixed parameters as in our final experimental design. Over these 23 sessions, performance had plateaued, with a median ARAT score of 18 and an interquartile range (IQR) of 16.25 – 19 (Fig. 3a). We then began collecting data to compare the effect of ICMS on ARAT performance. In the first block of four sequential sessions–which included ICMS, enabling our participant to feel tactile sensations perceived as originating from his own hand when the robotic hand grasped an object–his ARAT score increased significantly to a median of 21 and a range of 20 – 21 (U = 5, P = 0.005, Wilcoxon rank-sum test, Table 1, Fig. 3a). Performance with ICMS was also significantly better than the four subsequent matched control sessions without ICMS in which he achieved a median ARAT score of 17 with a range of 16 – 19 (U = 0, P = 0.029, Wilcoxon rank-sum test, Fig. 3a). ARAT scores in these control sessions were no different than the 23 historical sessions (U = 39, P = 0.65, Wilcoxon rank-sum test, Fig. 3a). Individual session scores are shown in Table 1. Despite the significantly improved scores in sessions with ICMS, there was no change in the total number of trials that were successfully completed (U = 7, P = 0.83, Wilcoxon rank-sum, Table 1). Therefore, the improved ARAT scores occurred as a result of completing individual trials more quickly. In the ARAT scoring system, successfully transferring an object in less than five seconds, and achieving a score of three, is considered normal, unimpaired performance^18^. In the absence of tactile sensations evoked by ICMS, a score of three was achieved only once during the 4 sessions (108 trials). When tactile sensations were provided, a score of three was attained 15 times during the 108 trials.

**Fig. 3:**
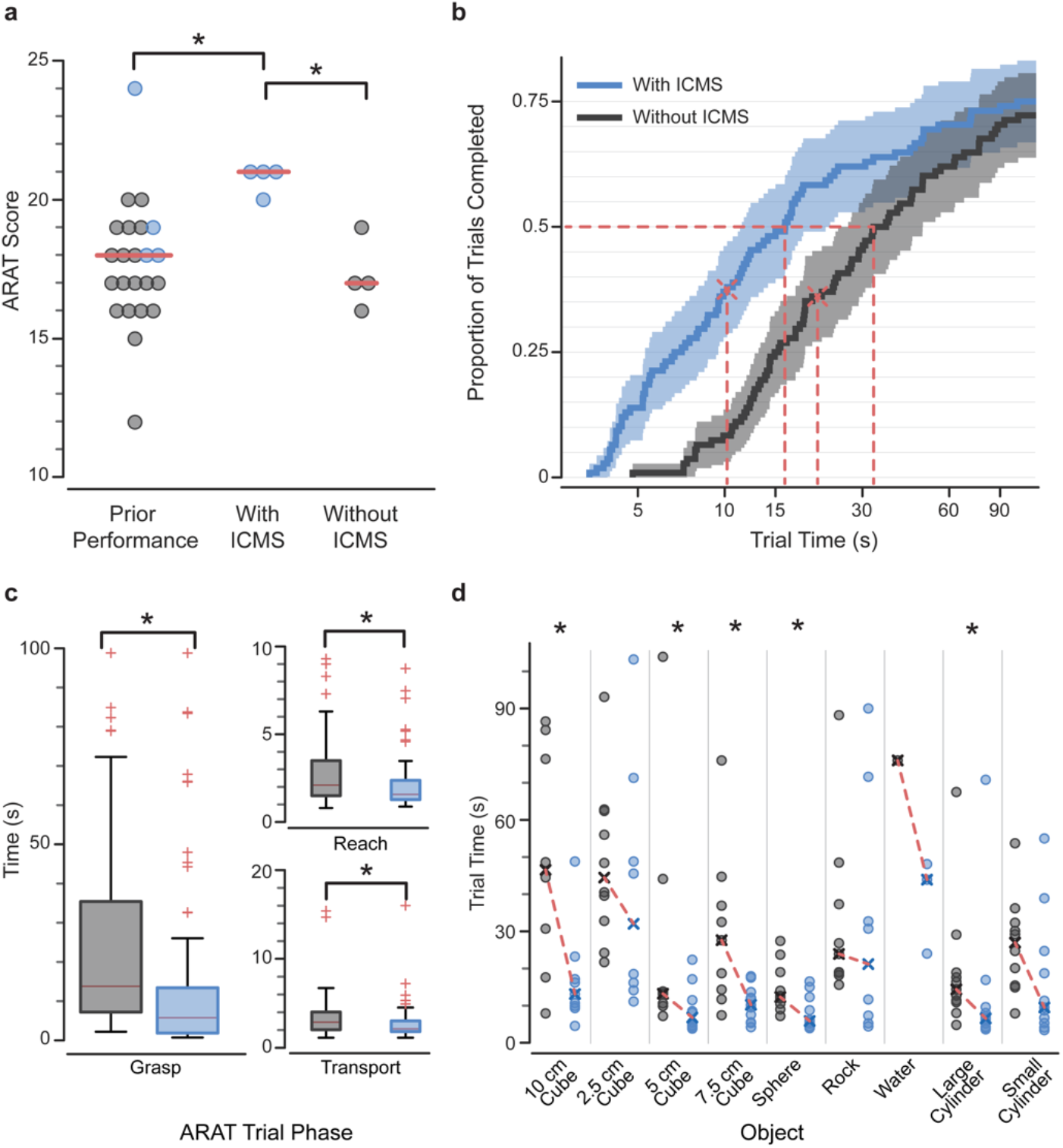
Effect of ICMS on ARAT task performance. **a**, Comparison of ARAT scores before experiment onset, which spanned a range of controlled degrees of freedom and occasionally employed ICMS feedback (blue dots), to data from the current experiment with ICMS feedback (blue) and without (gray). ARAT scores with ICMS feedback were significantly higher than historic performance (*P = 0.005, Wilcoxon rank-sum test) as well as control tests (*P = 0.029, Wilcoxon rank-sum test) conducted without ICMS. Red lines indicate the median score per paradigm. **b**, Cumulative distribution of individual trial times, including failed trials, shown on a log-normalized axis. Trials for all four sessions for each feedback condition were combined to compute the empirical cumulative distribution. The horizontal red line and connected vertical red lines indicate the times at which 50% of all attempted trials were completed for each condition. Vertical dashed lines connected to red X’s indicate when 50% of successful trials were completed. Shading indicates the 95% confidence bounds, calculated with Greenwood’s formula. **c**, Amount of time spent in each phase of the ARAT task. Red lines are medians, box outlines are interquartile ranges, and whiskers are the range of the data excluding outliers which are shown as red ‘+’ symbols. All task phases were faster when ICMS feedback was provided (*P < 0.01, Wilcoxon rank-sum test). For this analysis we included trials containing a successful reach, grasp and transport phase. Water pouring trials were not included as the transport phase is not defined. n = 78 trials for all phases without ICMS feedback and n = 85 trials for all phases with ICMS feedback. **d**, Effect of ICMS feedback on completion times for individual objects. Gray dots indicate trial times without ICMS feedback while blue dots are individual trial times with ICMS. Median trial times are marked for each object/feedback paradigm with an X. Medians for each object are connected with a red line for visualization. Trial times were significantly lower for five of the nine objects when ICMS feedback was provided (*P < 0.05, Wilcoxon rank-sum test).

Overall, we found that trials were consistently completed much more quickly when ICMS feedback was delivered (Fig. 3b, Supplemental Video 2); 14% of the trials with ICMS-evoked tactile feedback were completed more quickly than the fastest trial without ICMS. In fact, discounting the single trial that was completed in less than five seconds without ICMS, 25% of the trials with ICMS were completed more quickly than trials without ICMS (Fig. 3b). Successfully completed trial lengths decreased from a median time of 20.9 s (13.1 −40.5 s IQR) to 10.2 s (5.4 −18.1 s IQR) when tactile feedback was provided (U = 1676, P < 0.0001, Wilcoxon rank-sum test, Table 1, Fig. 3b and Supplemental Video 3). These faster completion times were the cause of the 3.5-point improvement in the ARAT score that occurred when ICMS was provided and can be interpreted as meaning that ICMS-induced tactile sensations allowed 3.5 more objects, out of 9 possible, to be transported to the platform in a normal time (< 5 seconds). The improved times were not due to differences in the commanded velocities. While the distributions of translation velocity commands measured at each time step were statistically different between conditions (D = 0.02, P < 0.0001, 2-sample Kolmogorov-Smirnov test, Extended Data Fig. 1), the velocities were functionally equivalent. The median translation velocity was 16.7 cm/s (11.5 – 23.2 cm/s IQR) with ICMS and 16.4 cm/s (11.4 – 22.6 cm/s IQR) without ICMS. Similar results were observed for wrist rotation and grasp velocities (Extended Data Fig. 1).

The ARAT task can be broadly divided into reach, grasp, and transport phases (Supplemental Video 4). We separated the trials into these three sequential task phases: (1) reach, consisting of movement onset to first object contact; (2) grasp, consisting of first object contact to successful object liftoff; and (3) transport, consisting of object liftoff to object release. The median time spent reaching decreased from 2.1 s (1.5 – 3.5 s IQR) without ICMS to 1.5 s (1.2 – 2.3 s IQR) when ICMS was provided, representing a 27.8% improvement (n = 78 without ICMS and n = 85 with ICMS, U = 2204, P = 0.0002, Wilcoxon rank-sum test, Fig. 3c). Likewise, the median time spent transporting the object decreased from 2.9 s (2.0 – 4.0 s IQR) to 2.1 s (1.8 – 3.0 s IQR), representing a 22.3% improvement (n = 78 without ICMS and n = 85 with ICMS, U = 2366.5, P = 0.002, Wilcoxon rank-sum test, Fig. 3c). Most impressively, the amount of time spent attempting to grasp the object decreased from 13.8 s (7.2 – 35.4 s IQR) without ICMS to 5.8 s (1.9 – 13.5 s IQR) with ICMS, resulting in a 44.7% improvement in performance (n = 78 without ICMS and n = 85 with ICMS, U = 1819.5, P < 0.0001, Wilcoxon rank-sum test, Fig. 3c). We speculated that, much like in the object transfer task, the participant spent less time attempting to grasp the objects during trials with ICMS-evoked tactile percepts because the percepts increased his certainty about object contact timing and his confidence that he had successfully grasped the object. Why the amount of time spent in the other two phases decreased is less clear. Since object contact and contact force cannot be felt without ICMS, he may have taken longer positioning the hand to improve the amount of information about object interaction he could extract visually, thus increasing the amount of time spent reaching. For the transport phase, the participant may have been less confident about his grasp stability, causing him to move more slowly during transport to avoid dropping the object.

By design, the objects in the ARAT task vary in size, shape, weight and, therefore, the overall difficulty in grasping them. As a result of the significant time spent practicing this task, the participant had classified the nine ARAT objects as being either easy (5 cm cube, 7.5 cm cube and sphere) or difficult (2.5 cm cube, 10 cm cube, rock, small cylinder, large cylinder and water pouring) to complete. All of the objects that were rated as easy, as well as the 10 cm cube and large cylinder, were completed more quickly with ICMS than without ICMS (Fig. 3d, Extended Data Table 1). Including ICMS did not significantly improve perfomance with the rock, small cylinder or water pouring task although the median completion time did go down for all of the objects. Therefore, other factors, such as the controllable degrees of freedom or kinematic constraints in the robotic arm, may have limited performance on these objects. However, for those objects that could be completed more easily, adding ICMS feedback further improved performance.

Prior to conducting the functional tasks each session, BCI decoder performance was tested in the absence of ICMS-evoked tactile feedback using a random target sequence task^22^. This task explicitly measured how well the participant could independently control each DoF by moving to specific locations in the 5 DoF workspace. On the days when ICMS-evoked tactile feedback was not provided, sequence task performance was slightly higher, achieving a median score of 100% on all four days compared to a median of 95% (range 90-100%) on the days where ICMS was delivered during the functional tasks (12 scores per condition, U = 40.5, P = 0.025, Wilcoxon rank-sum test, median scores for individual sessions in Table 1). This suggests that decoder performance itself, and thus the participant’s ability to control the robotic arm, did not favor the days on which ICMS was provided.

In many bidirectional upper-limb prosthetics studies where amputees receive restored sensory feedback through electrical stimulation of the peripheral nerves, the effect of artificial sensations on performance are measured without visual or auditory feedback^12,23–25^. Our approach differed from these studies in that our aim was to investigate the effect of providing artificial somatosensory feedback on tasks that were already possible with existing sensory modalities, namely vision. Here, we demonstrated that in highly-practiced tasks where normal visual feedback was available, adding artificial tactile feedback through ICMS enabled a person with spinal cord injury using a BCI to significantly improve their task scores, primarily by spending less time attempting to grasp the objects (Fig. 2a,c, 3b,c).

As with any single-subject study, it is uncertain whether these findings will generalize to future experiments. However, there are several reasons to believe that these results accurately represent the potential of restoring somatosensory percepts using ICMS. First, using the same fundamental neural decoding and control methods, we have demonstrated that two participants achieved similar scores on functional tasks with vision alone^20,22^ and that these scores were only exceeded when ICMS-evoked tactile feedback was provided (Fig. 3a). This suggests that without artificial tactile feedback, control is impaired, much as it is when tactile sensations are absent in people with otherwise normal motor control capabilities^3,26^. Second, we found that performance improvements were driven primarily by reductions in the time taken to successfully grasp an object. State transitions, such as object contact^5^ during the grasp phase, are uniquely encoded by tactile feedback in the intact nervous system. That the percepts signaled these state transitions with high temporal accuracy, and enabled him to grasp objects more quickly, suggests that ICMS delivered to area 1 of S1 can improve task performance in a way that is congruent to the way natural cutaneous feedback improves grasp performance. Finally, when ICMS-induced percepts were provided, performance improved significantly, and when they were removed, performance returned to pre-ICMS levels (Fig. 3a). Therefore, these observations suggest that the observed improvements were primarily due to the addition of reliable sensory information, rather than the result of additional practice. This immediate performance improvement also suggests that ICMS in S1 was not akin to sensory substitution cues that could have been provided by electrical or mechanical stimulation of intact skin or audio or visual cues, as the relationship between these cues and behavior must be learned^27^.

Ultimately, ICMS-induced tactile percepts improved task performance to levels not previously observed, decreased the time spent grasping in ways that were analogous to the role of natural tactile sensations during grasp state transitions, and do not appear to be the result of practice, suggesting that including naturalistic somatosensory feedback, like that induced with ICMS, could have a major impact on the future development and performance of dexterous prosthetic limb systems.

## Methods

### Implantation and electrode arrays

This study was conducted under an Investigational Device Exemption from the U.S. Food and Drug Administration and is registered at ClinicalTrials.gov (NCT01894802). The study was approved by the Institutional Review Boards at the University of Pittsburgh and the Space and Naval Warfare Systems Center Pacific. Informed consent was obtained before any study procedures were conducted.

A 28-year-old male participant with tetraplegia due to a C5 motor/C6 sensory ASIA B spinal cord injury was implanted with two sets of microelectrode arrays (Blackrock Microsystems, Inc., Salt Lake City, Utah, Fig. 1b). Two intracortical microelectrode arrays with 88 wired channels (10×10 array, 1.5 mm length platinum electrodes) were implanted in the hand and arm region of M1 in order to decode movement intent. Two additional microelectrode arrays with 32 wired channels were implanted in area 1 of S1 (6×10 array, 1.5 mm length and coated with a sputtered iridium oxide film) in order to evoke sensations in the fingers of the right hand when stimulated^7^. The study sessions described here took place between 717 and 738 days after the arrays were implanted.

### Neural Recording

Voltage recordings from each electrode were band-pass filtered between 0.3 Hz and 7.5 kHz and digitized at 30,000 samples per second using a NeuroPort signal processor (Blackrock Microsystems, Inc., Salt Lake City, Utah). Electrical artifacts induced by microstimulation were rejected using a combination of digital signal blanking and filtering. During each stimulus pulse the recorded signals were blanked using a sample-and-hold circuit. The signals were then high-pass filtered using a 750 Hz first-order Butterworth filter that minimized the effect of additional transient discontinuities in the signal, enabling fast settling of the wideband signal to baseline. A spike threshold was set at −4.5 times the root-mean-square of this high-pass filtered signal. Any transient threshold crossings that occurred in the sample immediately after the blanking period were rejected in software. Using this approach, we were able to record single unit activity within 740 μs of the end of a stimulus pulse^28^.

### Motor decoding

To investigate the ability of the participant to use ICMS-evoked tactile percepts during continuous control of a prosthesis, we first created a mapping between population-level neural firing rates recorded in M1 and desired arm movements. A 5 DoF decoder was used in this study, comprising translation of the endpoint in 3D space, wrist pronation and supination, and flexion and extension of all fingers and the thumb, with the thumb always opposite the fingers. All 5 DoFs were controlled simultaneously. A 5 DoF control scheme was chosen as it provided a balance between fast training times and a sufficient degree of dexterity to grasp the different objects used in these experiments.

To train the decoder, the participant observed a virtual version of the Modular Prosthetic Limb (MPL)^19^ moving in a 3D environment, as has been described previously^20^. In this task, the participant was asked to observe and imagine performing the motions of the MPL as the hand was first translated, then oriented, and finally commanded to grasp targets that were randomly presented throughout the workspace using a combination of virtual objects and auditory cues. After observing the completion of 27 trials, which took approximately 7 minutes, an optimal linear estimator decoder was derived using an encoding model that relates neural firing rates to arm kinematics. The encoding model was:

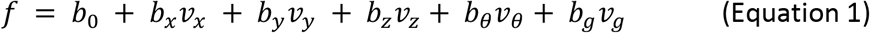

where *f* is the square-root transformed firing rate of a recorded unit, *v* is a kinematic velocity, and *b* is a regression coefficient for a given velocity dimension. The dimensions shown in Equation 1 are *x*, *y*, and *z* translation, wrist rotation (*θ*), and grasp (*g*). The *b* coefficients were calculated using linear regression^29^. Decoder weights were then calculated using indirect optimal linear estimation (Fig. 1e)^30^.

The participant used the decoder trained from observation data to repeat the training task, however the computer constrained the decoded movement velocities to those that were on the ideal path^31^. Once this task was completed, a new decoder was trained using the data from the second training set. During task performance, all firing rates were scaled, prior to being decoded, by dividing them by the ratio between the population firing rate during the most recent 300 ms and the population firing rate during decoder calibration. This method of scaling firing rates prior to decoding was developed to compensate for a correlated increase in firing rate across the recorded population that we observe when the prosthetic hand approaches objects^32^. This scaling allowed the participant to better stabilize the hand near objects in order to grasp them. Ultimately, this velocity decoder was then used, without computer assistance–that is the decoders and prosthetic arm control systems were naïve to the goal–to complete the tasks used to evaluate performance.

Decoder performance was evaluated using the physical MPL in a sequence task, where the goal was to acquire instructed combinations of hand endpoint position, wrist orientation and grasp posture^20,22^. A total of 3 sets of 10 trials were performed with the robotic limb without computer assistance to establish the baseline decoder performance accuracy in the absence of objects and ICMS. A trial was considered successful if the participant was able to place the robotic hand within a position target that was 5 cm in diameter, orient the wrist to within ± 0.25 radians and control the grasp aperture to be at least 80% of the way to maximum flexion or extension of the digits being used.

### Intracortical microstimulation

Stimulation pulse trains consisted of cathodal phase first, current-controlled, charge-balanced pulses delivered at a rate of 100 pulses per second. The cathodal phase was 200 µs long, the anodal phase was 400 µs long, and the amplitude of the anodal phase was set to half the amplitude of the cathodal phase. The phases were separated by a 100-µs interphase period. Detailed descriptions of sensory percepts evoked via ICMS of S1 have previously been reported^7^. Briefly, ICMS elicited percepts that were described by the participant as originating from the bases of the 2^nd^ through 5^th^ digits and up to the distal interphalangeal joint of the index finger. We selected the electrodes used to provide ICMS-evoked tactile percepts prior to the experiments and focused on electrodes that elicited easily detectable percepts with a clear projected location. One electrode, with a projected field in the proximal interphalangeal joint of the index finger, was mapped to the output of the torque sensor located at the index finger metacarpal phalangeal joint of the MPL. Four electrodes with projected fields in either the middle, ring or little finger were mapped to the torque sensor output from the middle finger of the MPL (Fig. 1c). Together, the projected fields from the selected electrodes spanned the index, middle, ring and little fingers.

For tasks with ICMS, torque sensors located in the motors controlling the MPL fingers provided the signal that was used to modulate ICMS pulse train amplitude according to the follow equation:

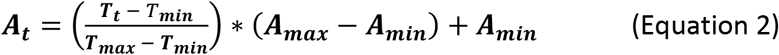

where *A*_*t*_ refers to the commanded pulse train amplitude at time step *t*, *A*_*min*_ and *A*_*max*_ refer to the electrode-specific range of stimulus amplitudes, and *T* represents the torque sensor data that was being used to relay grasp force. We also set values for the minimum and maximum torque readings, *T*_*min*_ and *T*_*max*_, respectively, that corresponded to the minimum and maximum stimulation amplitudes. The selected torque thresholds were 0.1 Nm and 0.5 Nm, which corresponded approximately to light touch and strong grasp, respectively. These values were linearly mapped to stimulus amplitudes that ranged from 14 to 64 µA in increments of 4 or 6 µA (Fig. 1d). New torque values were sampled every 20 ms and used to update the pulse train amplitude in real time.

### Functional task descriptions and scoring metrics

We used two different paradigms to quantify the effects of providing ICMS on the participant’s ability to complete functionally relevant tasks. Both the object transfer task and Action Research Arm Test (ARAT) have been successfully performed with vision as the only source of feedback^20,22^. Here we directly compared performance with and without ICMS-evoked tactile percepts while vision was always present.

For the object transfer task, we asked the participant to reach to and grasp a cylindrical object (16 cm tall and 4.3 cm in diameter) with a weighted base placed on the left side of the table, lift the object off of the table, carry it to the target area on the right, and release the object (Fig. 1f). Two boundaries were marked on the table that defined a 22.5 cm region where the object was not allowed to touch the table (red area in Fig. 1f). If the object touched the table between these boundaries, the task could be continued by moving the object back to the left side of the table and continuing. Once the object was placed on the right side of the table, an experimenter returned the object to the start position and the participant repeated the process as many times as possible in two minutes (Supplemental Video 1). Performance on this task was measured as the number of times the object was successfully moved across the table in two minutes. This task was always completed prior to the ARAT task.

We also conducted a modified version of the Action Research Arm Test (ARAT)^18,33^, which consisted of moving eight different objects from the left side of a table to a raised platform located on the right side (Fig. 1g). These objects were selected from the suite of objects that are part of the standard ARAT task^33^ and included four cubes (2.5 cm, 5 cm, 7.5 cm and 10 cm along each edge), a 7.5 cm diameter ball, a rock, and two cylinders (2.5 cm and 1 cm in diameter and 16 cm tall). Additional objects from the ARAT task were too small to be grasped by the MPL. The target platform was 34 x 20.5 cm and was elevated 12 cm off the table surface. The objects started approximately 70 cm away from the target platform. A ninth object from the original ARAT task was also included in which a cup filled with small pieces of paper and plastic, as a proxy for water, was placed at the right side of the workspace, and an empty cup was placed 20 cm to the left of it. The participant’s task was to empty the “water” from the cup on the right into the empty cup on the left and replace the originally grasped cup back on the table in an upright position. This task was considered a success if any “water” landed in the target cup and if the original cup was placed upright on the table.

In all cases, the participant was instructed to complete the task as quickly as possible. The participant had a maximum of two minutes per attempt, and three attempts per object. Each attempt at transferring the objects was considered a trial. Trials were timed by experimenters from movement onset to the object being successfully placed on the target platform. Each trial was scored on a 3-point system in which a score of zero was awarded if the object was never touched, a score of one was awarded if the object was touched but the participant was unable to complete the task, a score of two was awarded if the task was completed in less than two minutes but more than five seconds, and a score of three was awarded if the task was completed in under five seconds. The best score from the three attempts for each object was added together to create a single score for the test. Therefore, for the task with nine objects, a perfect score was 27.

The score, which is the validated metric of the ARAT task, fails to take into account other aspects of performance, such as the total number of completed attempts per object and the actual completion time. Therefore, we recorded video of all trials and measured the time spent reaching for, grasping, and transporting the object. All task phase calculations were done offline, marking individual video frames that spanned each event. Reaching was defined as the time from movement onset until the first object contact. Grasping was defined as the period between object contact and successful object liftoff from the table. The transport phase spanned object liftoff until object release.

We tested the two feedback conditions in a block-design over the course of these experiments. For the first four sessions, ICMS feedback was delivered to five electrodes. Each experiment day, three blocks of the sequence task, five blocks of the object transfer task, and one ARAT session were completed. For the next four consecutive sessions, the same testing protocol was followed, but ICMS was not delivered.

### Statistical analysis

Statistical analyses were performed in MATLAB (The MathWorks). Data that were not normally distributed, as determined using Lilliefors test (α = 0.05), are reported as medians and interquartile ranges (IQR) and the Wilcoxon rank-sum test was used to assess significance for differences in the median unless otherwise stated. The Mann-Whitney U test statistic is reported for all Wilcoxon rank-sum tests. Normally-distributed data, as determined using Lillifors test (α = 0.05), are reported as mean ± standard deviation and a two-tailed Student’s t-test was used to assess significance for differences in the mean. Specific statistical tests are noted in the text. All object transfer data have n = 20 trials per feedback condition.

### Data availability

Data supporting these findings as well as software routines to analyze these data are available from the corresponding author upon reasonable request.

## End Notes

### Supplementary Information

is available in the online version of this paper.

## Acknowledgements

We thank N. Copeland for his continuing and extraordinary commitment to this study as well as insightful discussions with the study team; Debbie Harrington (Physical Medicine and Rehabilitation) for regulatory management of the study; Ahmed Jorge for help with data collection; Peter Gibson and Ben Clarkson for video data processing; the University of Pittsburgh Clinical and Translational Science Institute and the Office of Investigator-Sponsored Investigational New Drugs and Investigational Device Exemption support for assistance with protocol development and regulatory reporting and compliance; the volunteer members of the Data Safety and Monitoring Board for their continued monitoring of this study; H. Jourdan (Department of Physical Medicine and Rehabilitation) for financial and organizational support; and Blackrock Microsystems (Salt Lake City, UT, USA), especially Robert Franklin, for technical support related to this project. This material is based upon work supported by the Defense Advanced Research Projects Agency (DARPA) and Space and Naval Warfare Systems Center Pacific (SSC Pacific) under Contract No. N66001-16-C-4051 and the Revolutionizing Prosthetics program (Contract No. N66001-10-C-4056). S.N.F. was supported by an NSF Graduate Research Fellowship under grant number DGE-1247842. The views, opinions, and/or findings contained in this article are those of the authors and should not be interpreted as representing the official views or policies of the Department of Veterans Affairs, Department of Defense, or US Government.

## Author Contributions

S.N.F., J.E.D., J.L.C., and R.A.G. designed the study. S.N.F., J.E.D., J.M.W., C.L.H., A.J.H., J.L.C., and R.A.G. conducted the experiments. S.N.F. analyzed the data. All authors contributed to the interpretation of the results. S.N.F. wrote the paper with R.A.G. and J.L.C., and all authors provided critical review, edits, and approval for the final manuscript.

## Author Information

The authors declare that they have no competing interests. Correspondence and requests for materials should be addressed to R.A.G and J.L.C.

## Extended Data

**Extended Data Fig. 1:**
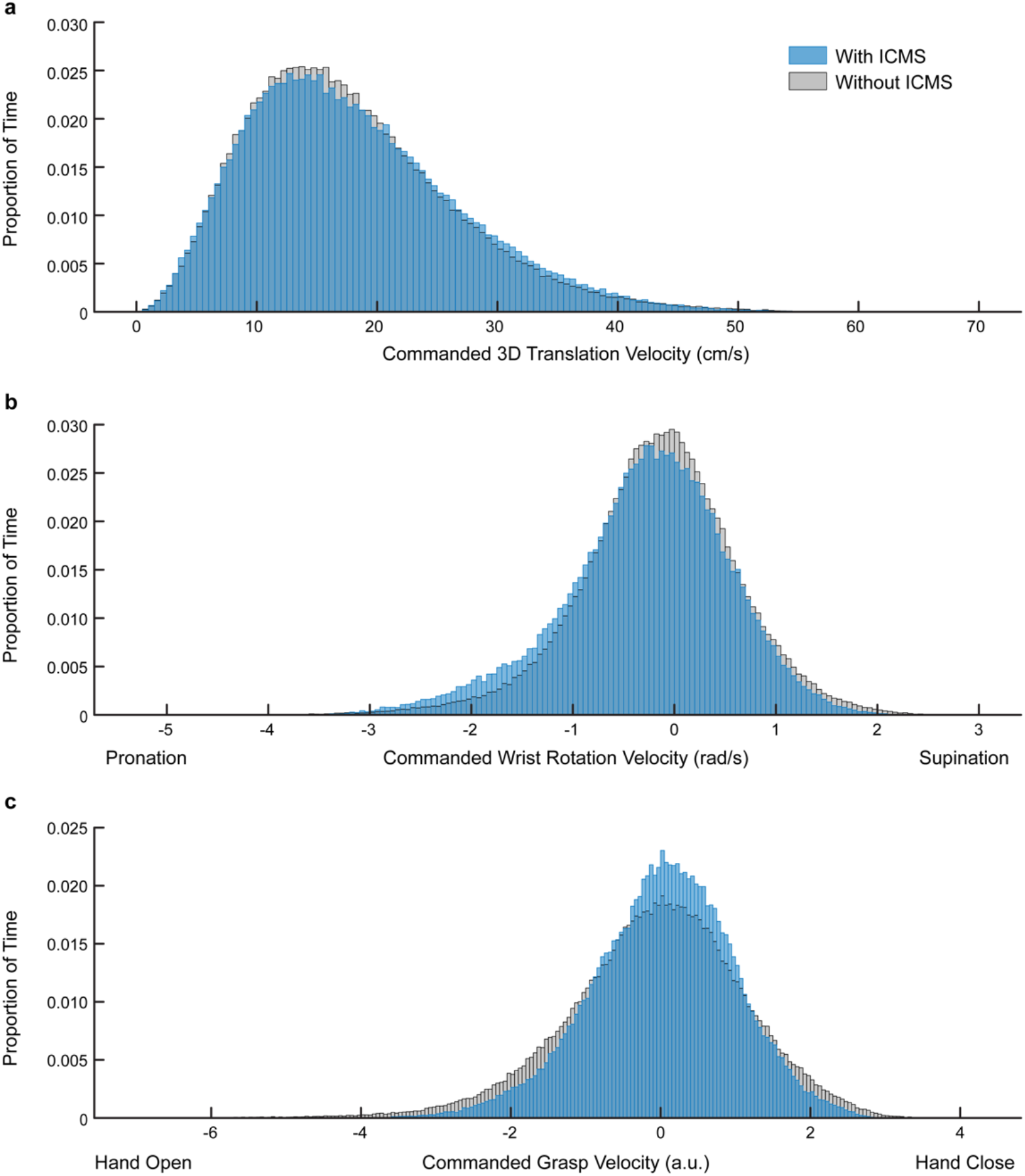
Distribution of commanded robot velocities for each timestep during all ARAT trials with (blue) and without ICMS (gray) **a**, Commanded 3D translation velocity. The distribution of commanded translation velocities were different for trials with and without ICMS (D = 0.02, P < 0.001, 2-sample Kolmogorov-Smirnov test). **b**, Commanded wrist rotation velocity. The distributions of commanded wrist rotation velocities were different for trials with and without ICMS (D = 0.055, P < 0.0001, Kolmogorov-Smirnov test). The median wrist rotation velocity was −0.22 rad/s (−0.74 – 0.26 rad/s IQR) with ICMS and −0.13 rad/s (−0.61 – 0.33 rad/s IQR) without ICMS. **c**, Commanded grasp velocity. The distributions of commanded grasp velocities were different for trials with and without ICMS (D = 0.058, P < 0.0001, Kolmogorov-Smirnov test). The median grasp velocity was 0.074 a.u. (−0.571 – 0.680 a.u. IQR) with ICMS and −0.001 a.u. (−0.763 – 0.711 a.u. IQR) without ICMS.

**Extended Data Table 1:** Successful ARAT trial times by object. All successful water pouring attempts are listed as there were not enough successfully completed trials to calculate the median and IQR.

| Object | Median (s) | IQR (s) | n | Mann-Whitney<br>U Statistic | Significance (p-value, Wilcoxon<br>rank-sum test) |
| --- | --- | --- | --- | --- | --- |
| 10 cm cube – no ICMS | 46.6 | 24.2 – 80.3 | 8 | 13 | 0.027 |
| 10 cm cube – ICMS | 13.1 | 9.8 – 18.6 | 9 |  |  |
| 2.5 cm cube – no ICMS | 44.5 | 32.8 – 62.4 | 10 | 30 | 0.408 |
| 2.5 cm cube – ICMS | 32.0 | 15.1 – 60.1 | 8 |  |  |
| 5 cm cube – no ICMS | 13.2 | 10.3 – 29.0 | 8 | 17 | 0.043 |
| 5 cm cube – ICMS | 6.8 | 4.3 – 11.5 | 10 |  |  |
| 7.5 cm cube – no ICMS | 27.6 | 13.7 – 38.9 | 9 | 15 | 0.010 |
| 7.5 cm cube – ICMS | 10.2 | 6.0 – 13.3 | 11 |  |  |
| Sphere – no ICMS | 12.3 | 10.9 – 17.8 | 11 | 22.5 | 0.024 |
| Sphere – ICMS | 5.9 | 4.4 – 12.3 | 10 |  |  |
| Rock – no ICMS | 23.9 | 18.7 – 40.1 | 9 | 29 | 0.541 |
| Rock – ICMS | 21.2 | 6.3 – 52.2 | 8 |  |  |
| Large Cylinder – no ICMS | 14.4 | 11.2 – 18.3 | 12 | 27.5 | 0.019 |
| Large Cylinder – ICMS | 6.6 | 4.5 – 9.2 | 11 |  |  |
| Small Cylinder – no ICMS | 27.0 | 15.4 – 32.3 | 10 | 29.5 | 0.078 |
| Small Cylinder – ICMS | 9.5 | 5.7 – 23.2 | 11 |  |  |
| Water – no ICMS (all times) | 76 |  |  | n/a |  |
| Water – ICMS (all times) | 24, 43.9, 48.1 |  |  |  |  |

## Supplementary Information

**Supplemental Video 1:** Object transfer example trials with and without ICMS feedback. In the full trial, the task lasts for two minutes. The first minute from a trial with the median number of transfers for each feedback condition is used to illustrate performance.

**Supplemental Video 2:** Fastest ARAT trials for each object and feedback condition.

**Supplemental Video 3:** ARAT trials for the median completion time for each object and feedback condition. In cases where there were an even number of completed trials, the faster trial is shown in the video.

**Supplemental Video 4:** Example ARAT trial with ICMS feedback, annotated to indicate task state transitions and illustrate ICMS delivery.

